# Integration of a Computational Pipeline for Dynamic Inference of Gene Regulatory Networks in Single Cells

**DOI:** 10.1101/612952

**Authors:** Kyung Dae Ko, Stefania Dell’Orso, Aster H. Juan, Vittorio Sartorelli

## Abstract

Single-cell RNA-seq permits the characterization of the molecular expression states of individual cells. Several methods have been developed to spatially and temporally resolve individual cell populations. However, these methods are not always integrated and some of them are constrained by prior knowledge. Here, we present an integrated pipeline for inference of gene regulatory networks. The pipeline does not rely on prior knowledge, it improves inference accuracy by integrating signatures from different data dimensions and facilitates tracing variation of gene expression by visualizing gene-interacting patterns of co-expressed gene regulatory networks at distinct developmental stages.

## INTRODUCTION

The sequential and dynamic establishment and dismantling of gene regulatory networks (GRNs) instruct uncommitted and progenitor cells to adopt or avoid branching lineage choices (Davidson, 2006). Bulk transcriptomes have provided considerable insights and fostered the discovery and characterization of GRNs (Trapnell et al., 2010; Wang et al., 2009). However, bulk transcriptomes provide population-based averaged measurements which blur cell heterogeneity and developmental dynamics of asynchronous cell populations. Single-cell transcriptome technologies (scRNA-seq) capture cell heterogeneity and thus are useful for the discovery of cell populations, identification of cell mutants, and quantification of subpopulations (Linnarsson and Teichmann, 2016). Leveraging on the ability of generating thousands of individual measurements, methods have been developed to spatially and temporally resolve cell populations. Clustering and dimensionality reduction algorithms such as PCA, tSNE, and diffusion maps permit the identification and enumeration of cell types among cell populations (Satija et al., 2015) (Butler et al., 2018). Temporal trajectories are generated by pseudotime ordering of single cells to identify unique transition paths among different cell states (Trapnell et al., 2014) (Qiu et al., 2017) or by predicting future states of gene expression based on measurements of unspliced and spliced transcripts (La Manno et al., 2018). Using spatial or temporal information from clustering and trajectories, boolean models (Woodhouse et al., 2018), co-expression analysis (Allen et al., 2012), and multivariate information theory-based algorithms (Chan et al., 2017) have been successfully employed to infer GRNs. However, their accuracy depends on the size of the network (Fiers et al., 2018) and methods of normalization (Crow et al., 2016). To alleviate these issues, SCENIC (Aibar et al., 2017) combines co-expression with DNA binding motif enrichment analysis and SINCERA (Guo et al., 2015) makes use of scRNA-seq specific cell-type gene signatures. Because their power to infer GRNs is knowledge-based, the use of these models is constrained by the availability of annotated datasets (Fiers et al., 2018).

Here, we present an integrated pipeline for GRNs inference which uses clustering, temporal, and biological signatures extracted directly from scRNA-seq datasets. To evaluate its predictive power, we apply it to datasets derived from differentiating human pluripotent stem cells. The pipeline correctly identifies signaling pathways activated and dismantled at specific stages of cell differentiation and reveals the composition of gene hubs underlying discrete GRNs in pluripotent and committed human cells.

## RESULTS

### Pipeline Workflow

The <u>i</u>ntegrated co<u>m</u>putational pipeline for <u>s</u>ingle cell <u>ge</u>ne regulatory <u>n</u>etwork (IMSGEN) (Figure 1A) starts with the identification of transcripts corresponding to candidate genes in top 100 gene transcripts with average log fold changes among all cell clusters. Gene signatures are then employed to identify signaling pathways and gene ontology (GO) enrichment within each cluster. The clusters are temporally ordered by cell re-clustering using principal component analysis for dimension reduction and minimum spanning tree (MST) methods for trajectory modeling to predict the temporal relations among the clusters. After temporal ordering of cell clusters, distance matrices of dynamic and cluster-specific gene-interacting patterns are employed to infer GRNs by corrected gene interacting (CGI) maps and force-directed graph (FDG) network model (Fruchterman, 1991). This approach permits the identification and visualization of GRN transitions occurring in distinct cell states independent of cell-type and annotation biases.

**Figure 1.**
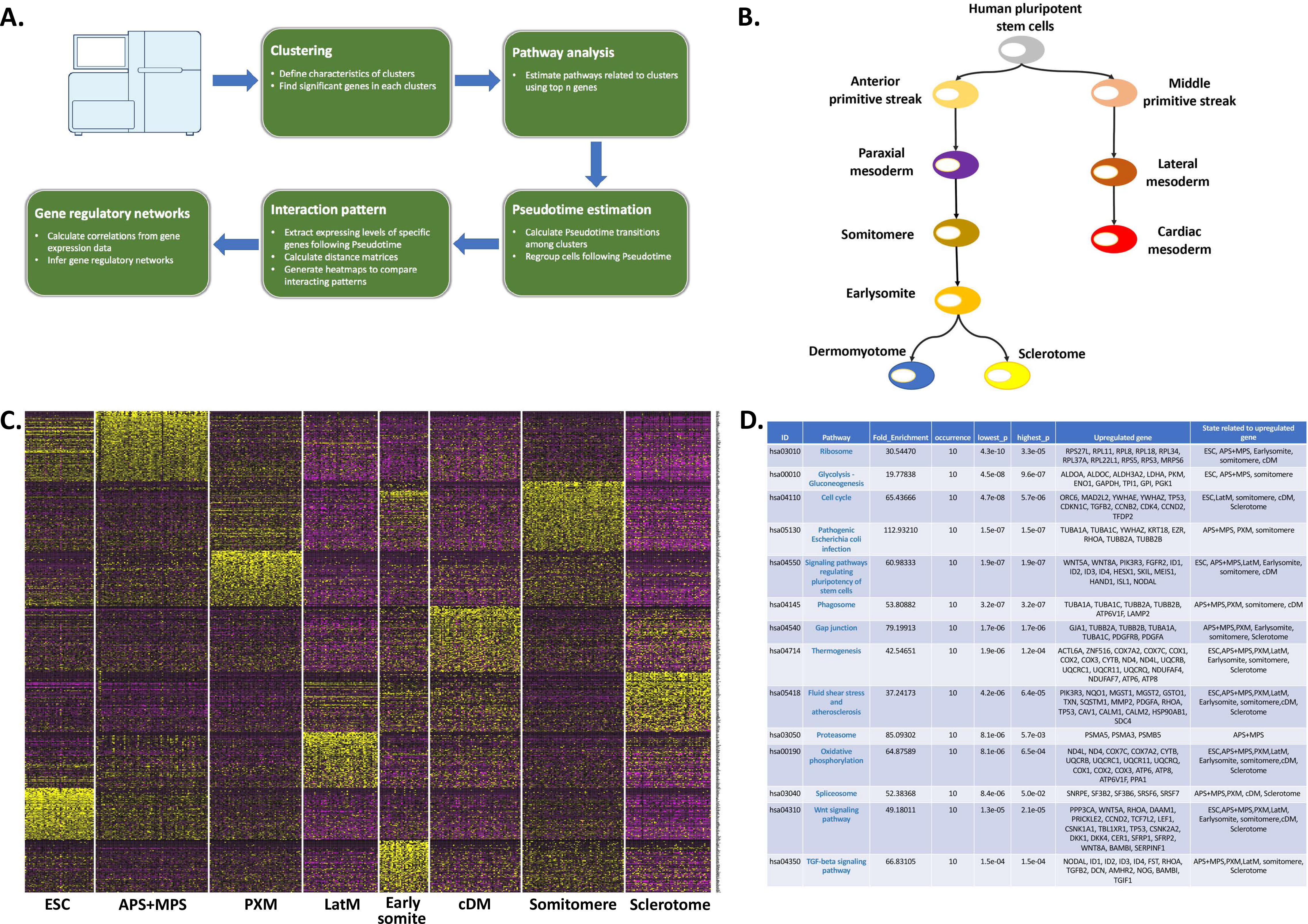
(A) Pipeline’s flowchart. (B) Scheme representing alternative mesoderm differentiation choices of human pluripotent stem cells. Left branch, somitogenesis; right branch, cardiogenesis (Loh et al., 2016). (C) Heatmap of top 100 most variably expressed genes in human pluripotent stem cells (ESC) and at the indicated stages of mesoderm differentiation. (D) Identification of pathways enriched in human pluripotent stem cells (ESC) and at the indicated stages of mesoderm differentiation.

### Clustering and Temporal Ordering of Human Pluripotent Cells undergoing Mesoderm Differentiation

We tested IMSGEN performance on experimental data generated from scRNA-seq of 498 human pluripotent cells undergoing mesoderm differentiation (Loh et al., 2016). scRNA-seq was performed on cells induced to differentiate and captured at specific developmental stages: 51 human pluripotent stem cells (embryonic stem cells, ESCs), cells differentiated into anterior and middle primitive streaks (59 APS and 22 MPS cells, respectively), paraxial mesoderm (67 PXM cells), lateral mesoderm (LatM 55 cells), somitomere (76 cells), early somites (36 cells), dermomyotome (67 cells), and sclerotome (65 cells) (Figure 1B). We aggregated scRNA-seq datasets without *a priori* knowledge of the developmental stage of the individual cells and clustered gene expression data using PCA and graph-based clustering method (Butler et al., 2018). Cells segregated into 8 clusters, representing each developmental stage with the exception of APS and MPS populations which could not be individually and correctly identified (Figure S1A). To evaluate somitogenesis (left branch of Figure 1B), we temporally ordered cell clusters by independent component analysis (ICA) for dimension reduction and MST for trajectory model (Figure S1B). Signatures for the individual clusters were generated by differential gene expression of the top 100 expressed genes (Figure 1C, Table S1) which were subsequently employed to perform pathway and GO enrichment analyses for the different clusters (Figure 1D, Figure S1C, Table S2). As expected, pathways regulating pluripotency of stem cells were identified and, consistent with their role in mesoderm induction (Cheung et al., 2012) (Gertow et al., 2013) (Loh et al., 2016), WNT and TGFβ pathways were also captured by this analysis. This unbiased approach permitted the identification of metabolic (glycolysis-gluconeogenesis) and several other pathways not directly queried in (Loh et al., 2016) (Figure 1D and see below).

### Identification of Signaling Pathways and Visualization of Gene Regulatory Networks

WNT and TGFβ pathways play key roles in mesoderm formation starting from human pluripotent stem cells (Cheung et al., 2012) (Gertow et al., 2013) (Loh et al., 2016). However, the composition, structure and temporal formation of WNT and TGFβ GRNs occurring at discrete differentiation states have not been elucidated. To infer GRNs in cells undergoing somitogenesis, we queried sub-datasets for WNT and TGFβ signaling pathways (Figure 1C,D and Table S1) and generated pairwise matrices to represent gene co-expression using normalized distance methods (Figure 2). In these matrices, gene proximity (red in Figure 2) indicates co-expression while gene distance (blue in Figure 2) indicates absence of co-expression, thus allowing to evaluate the presence of functionally connected and related modules (Stuart et al., 2003) (Nguyen and Lio, 2009). This analysis revealed formation of WNT GRN in ESC and APS (Figure 2A). In PXM and somitomere cells, the GRN’s strength was reduced and further decreased in early somite and sclerotome cells (Figure 2A). Similarly, gene interactions within a TGFβ GRN present in ESC and APS were diminished in PXM and somitomere cells and continued to decline in early somite and sclerotome cells (Figure 2B). These findings are consistent with an inhibitory role exerted by both WNT and TGFβ signaling pathways on early somitogenesis (Loh et al., 2016). Using the same approach, an embryonic morphogenesis GRN was revealed to be gradually established and to progressively increase its connectivity as cells progressed from pluripotent to more differentiated cell states (Figure 2C), reflecting morphogenesis that occurs during cell differentiation. Appropriate temporal expression of the somitomere-specific MESP2, HEYL, and HOPX, and somite-specific MEOX1 and PARAXIS (TCF15) and FOXC2 genes (Loh et al., 2016) confirmed that the matrices accurately represent the individual cell developmental stages (Figure S1D). Thus, using an unbiased approach, our computational pipeline correctly identified and ordered GRNs for pathways known to regulate human pluripotent cell differentiation. Converting similarity matrices to adjacency matrices, we visualized co-expression networks using FDG network model (Fruchterman, 1991). This way, we could identify the nodes (genes), edges (gene connectivity) and overall structures of the WNT, TGFβ, and embryonic morphogenesis GRNs at each different stages of somitogenesis (Figure S2A,B). Both WNT and TGFβ networks increased their connectivity during the transition from pluripotency (ESC) to APS. Genetic interactions were pruned and refined at later stages of somitogenesis (Figure 2A,B). In early somite cells, the transcription factors Smad2, TCF7L2 (TCF-4), the TCF-4 interacting corepressor CtBP1, and calcineurin (PPP3CA), known to be involved in mesoderm formation (Dunn et al., 2004) (Kardon et al., 2003) (Hogan et al., 2003), were found to establish a WNT subnetwork (Figure S2A). A TGFβ subnetwork revealed connectivity between the TGFβ-stimulated Rho-associated kinase ROCK1, important for somitogenesis (Wei et al., 2001) and the activin A receptor ACVR2B in early somite cells (Figure S2B). SMAD2 and the serine/threonine Protein Phosphatase 2 (PPP2R1A) were equally connected in both WNT and TGFβ subnetworks, confirming cross-talk of the two pathways (Attisano and Wrana, 2013). In contrast to the dismantling of the TGFβ and WNT GRNs, a GRN composed of genes related to embryogenic morphogenesis (Figure S2C) gradually increased connectivity acquiring additional nodes and edges in cells undergoing differentiation. Thus, IMSGEN was able to identify and dissect the composition, structure and temporal formation of GRNs in human pluripotent cells undergoing mesoderm differentiation.

**Figure 2.**
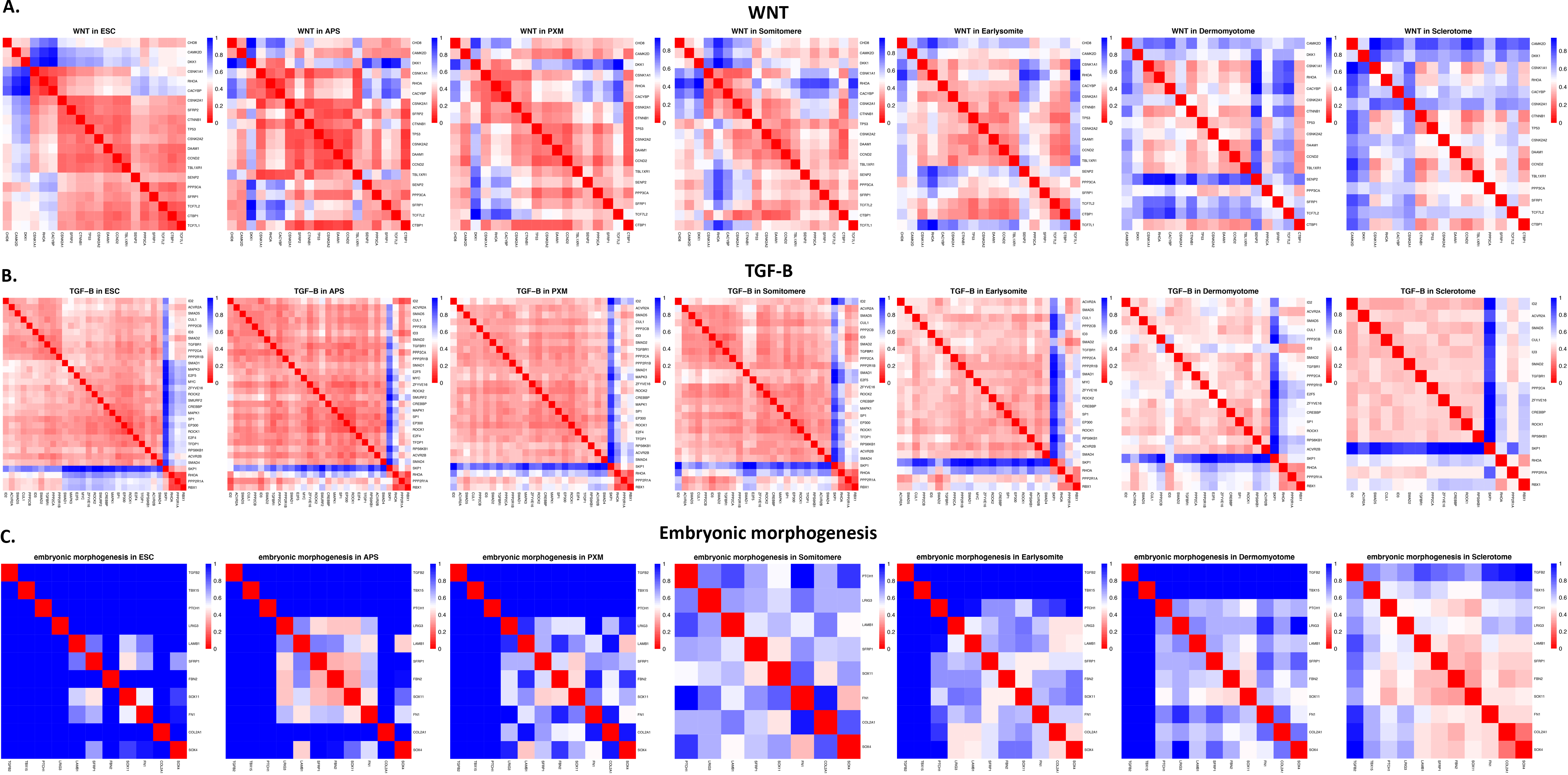
(A-C) Distance matrices indicating gene connectivity (highest connectivity, red; lowest connectivity, blue) for the WNT (A), TGFβ (B), and Embryonic morphogenesis (C) genes in human pluripotent stem cells (ESC) and at the indicated stages of mesoderm differentiation.

## DISCUSSION

Inference of GRNs from scRNA-seq data provides important clues to understand gene expression dynamics in developing systems. The pipeline described here, IMSGEN, complements and integrates existing methods. The salient characteristics of the pipeline are its independence from annotation biases, improved accuracy of inference integration of transcriptional signatures from different data dimensions, and easy visualization of gene interacting patterns and co-expressed GRNs. The pipeline performed well when tested with published data and its use can be extended to analyze GRNs during cellular development in any cell type and organism.

## ACKNOWLEGMENTS

We thank Dr. Hong-Wei Sun (Biodata Mining and Discovery Section, NIAMS) for critical reading of the manuscript. This work was supported by the Intramural Research Program of the National Institute of Arthritis, and Musculoskeletal and Skin diseases (NIAMS) at the National Institutes of Health (NIH grants AR041126 and AR041164).

## AUTHOR CONTRIBUTIONS

Conceptualization, K.D.K and V.S.; Software Design, K.D.K; Writing K.D.K, V.S.; Biological expertise and advice, S.D.O, A.H.J, V.S.

## WEB RESOURCES

GitHub, https://github.com/holyone09/lmsgen

Seurat, https://satijalab.org/seurat

Pathfind, https://cran.r-project.org/web/packages/pathfindR/index.html

Metascope, http://metascape.org/gp/index.html#/main/step1

Pheatmap, https://cran.r-project.org/web/packages/pheatmap/index.html

Igraph, https://igraph.org/redirect.html

## FIGURE LEGENDS

**Figure S1.**
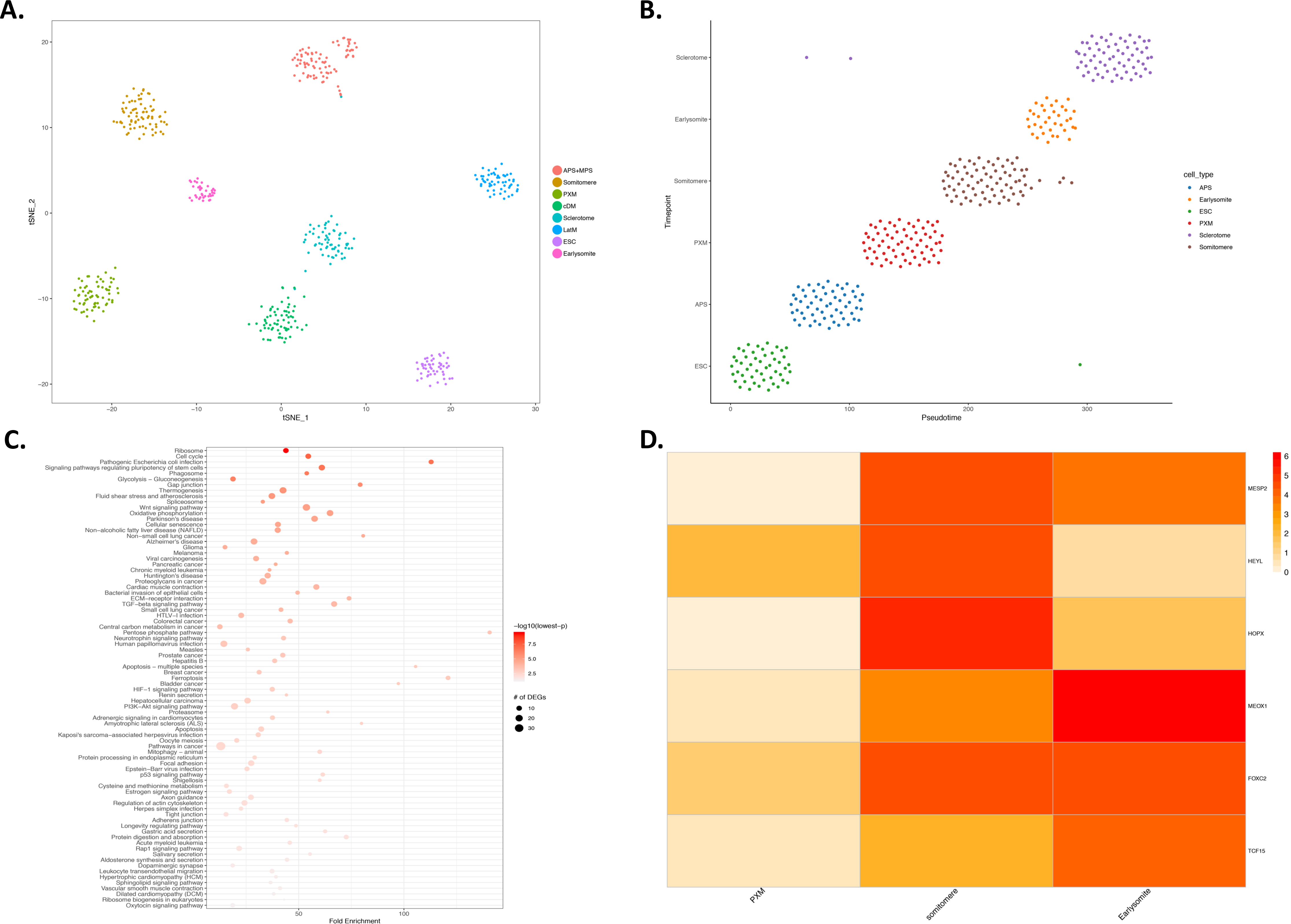
(A) tSNE-based clustering of human pluripotent stem cells (ESC) and of cells at different stages of mesoderm differentiation. (B) Pseudotime ordering of ESC and cells at different stages of somitogenesis. (C) Pathway enrichment representation based on pathways identified in Figure 1C. The size of the symbols for differentially-expressed genes (DEGs) is proportional to gene number (10,20, or 30 genes, respectively) and p-values of the single pathways indicated in the −log10 red color scale m (lower p-values, brighter red color). (D) Heatmap of MESP2, HEYL, HOPX, MEOX1, FOXC2 and TCF15 (PARAXIS) transcripts at different stages of mesoderm differentiation.

**Figure S2.**
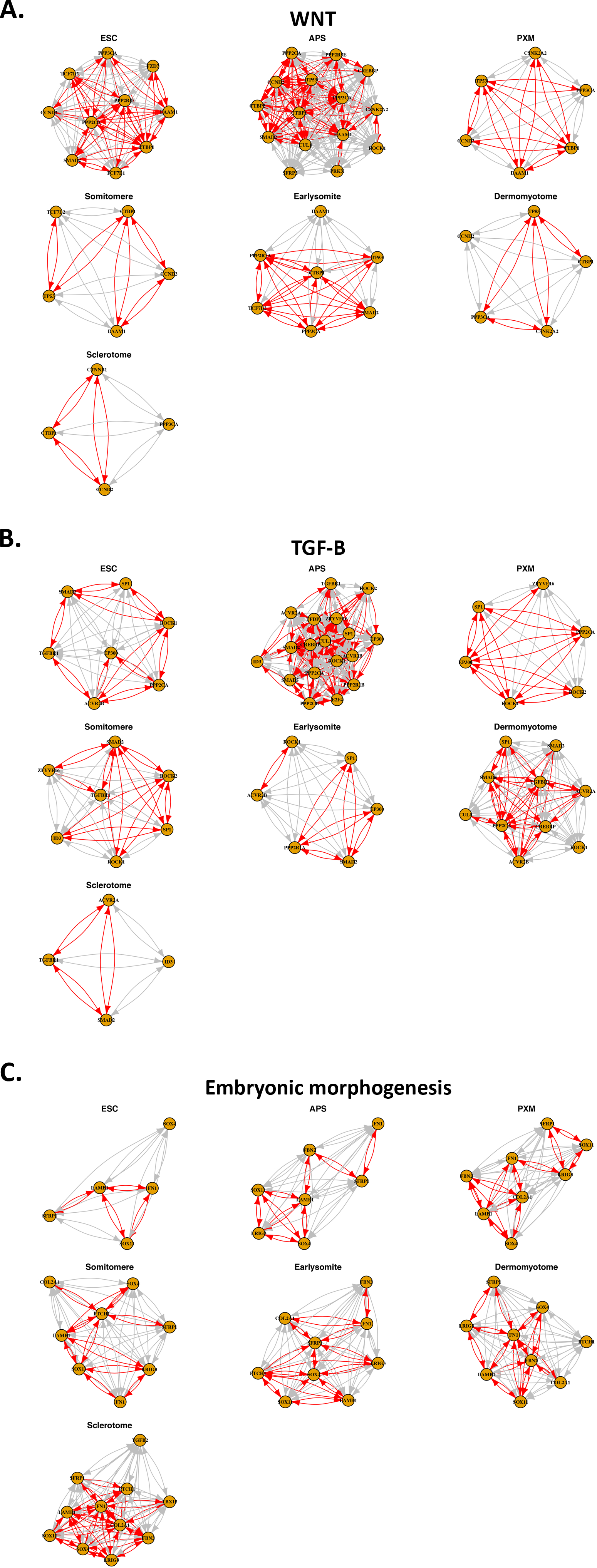
(A-C) Force-directed networks of WNT (A), TGFβ (B), and Embryonic morphogenesis (C) genes in human pluripotent stem cells (ESC) and at the indicated stages of mesoderm differentiation. Red edges indicate positive and grey edges negative gene correlation.

**Table S1.**List of the top 100 differentially expressed genes in human pluripotent stem cells (ESC) and in different stages of mesoderm differentiation.

**Table S2.**Pathways enriched in human pluripotent stem cells (ESC) and in different stages of mesoderm differentiation.

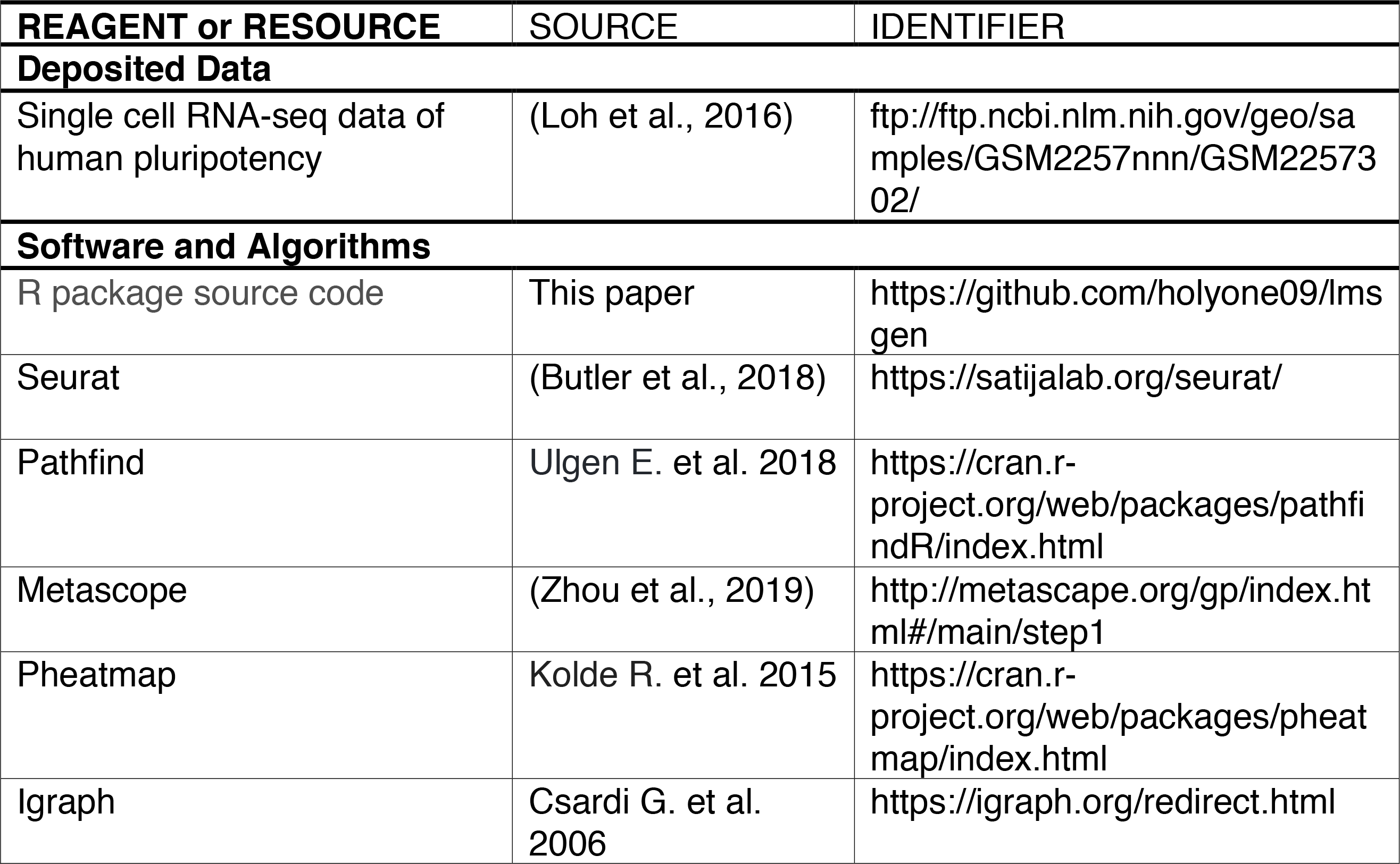
KEY RESOURCES TABLE.

### Contact for Reagent and Resource Sharing

Further information and requests for reagents should be directed to Lead Contact Vittorio Sartorelli.

### Method Details

The pipeline (Figure 1A) consist of five modules, and each module is implemented by R programming language with its package. To evaluate the effectiveness of analyzing results from the pipeline, we applied it to scRNA-seq datasets of human pluripotent stem cells (Loh et al., 2016).

### Extracting spatial, biological, and temporal signatures from datasets

The first step of the pipeline is to reduce the matrix of UMI counts or gene expressing values such as TPMs (Transcripts per millions) into PCA dimension. Based on Seurat R package (Butler et al., 2018), cells are clustered using graph-based clustering methods with PCA values and top-ranking genes are collected among clusters calculating average log fold changes. Next, the results of clustering and top-ranking gene list are transferred into the modules to identify biological signatures and reconstruct the temporal orders of clustered groups separately. Using Pathfind R package (Ulgen E. et al. 2018), we predict signaling pathways with low p-value (<0.01) for entire cell populations and gather candidate genes related to the pathways involved in cell differentiation. In addition, genes are extracted from GO (Gene Ontology) enrichment analysis (p < 0.01) of Metascope (Zhou et al., 2019). Finally, lists of genes are preprocessed to visualize gene interaction. To reconstruct temporal orders of clusters, we reduce the dimension of the distance matrix to PCA after calculating euclidean distances among scRNAseq data. Then, temporal signatures are restored from PCA by Mclust and MST (Minimum Spanning Tree) algorithms (Xu et al., 2002). Finally, we reorganize spatial clusters following temporal signatures.

### Visualization of gene-interacting patterns and GRNs

Using pre-processed gene lists, we generate the temporal submatrices of selected gene expression from an original expressing matrix. To visualize the patterns of gene interactions in the temporal submatrices, we generate the matrices of similarity calculating normalized distance methods with eq.(1), and we draw correlational heatmaps using the matrices of similarity using Pheatmap R package (Kolde R. et al. 2015).

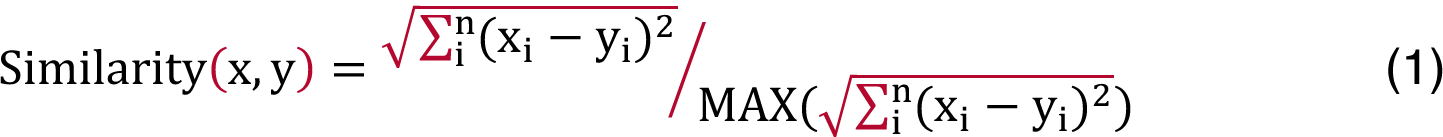

Converting these similarity matrices to adjacency matrices for co-expression networks, we visualize co-expression networks in each state using Igraph R package (Csardi G. et al. 2006) and the Fruchterman-Reingold layout algorithm (Fruchterman, 1991). Genes with concordant expression levels are closely positioned forming multiple hubs within a given GRN.

### Data and Software Availability

Source code and installation of the R package is available at https://github.com/holyone09/lmsgen under Open-source R package under ‘GPL (version 2 or later)’.

